# Drivers of avian genomic change revealed by evolutionary rate decomposition

**DOI:** 10.1101/2025.02.19.638997

**Authors:** David A. Duchêne, Al-Aabid Chowdhury, Jingyi Yang, Maider Iglesias-Carrasco, Josefin Stiller, Shaohong Feng, M. Thomas P. Gilbert, Guojie Zhang, Joseph A. Tobias, Simon Y. W. Ho

## Abstract

Modern bird diversity spans a striking array of forms, behaviours, and ecological roles. Analyses of molecular evolutionary rates can reveal the links between genomic and phenotypic change, but disentangling the drivers of rate variation has been difficult across large numbers of whole-genomes. Using comprehensive estimates of traits and evolutionary rates across a family-level phylogeny of birds, we show that clutch size, generation length, and beak shape are dominant predictors of genome-wide mutation rates. To identify the major axes of evolutionary rate variation, we employ covariance matrix eigendecomposition from rates estimated for branches of the avian phylogeny and across genomic loci. We find that the majority of rate variation occurs along the terminal branches of the phylogeny associated with extant families of birds. Additionally, we use principal components analyses to show that several axes of variation are linked with rapid evolution in microchromosomes immediately after the Cretaceous–Palaeogene transition. These apparent pulses of evolution are consistent with major changes in evolutionary rates in the machineries for meiosis, heart performance, and RNA splicing, surveillance, and translation. They also correlate with the diversification of ecological niches reflected in increased tarsus length. Collectively, our analyses paint a nuanced picture of avian evolutionary rates through time, revealing that the ancestors of the most diverse lineages of birds underwent major genomic changes related to mutation, gene usage, and niche expansion near the beginning of the Palaeogene period.

## Main

Birds are the most species-rich lineage of tetrapods and occur in all major habitat types on the planet. Their evolutionary radiation is associated with striking biological innovation in adaptive traits^1^, including flight^2,3^, song^4–6^, colouration^7,8^, and beak morphology^9,10^. These traits have various associations with genomic evolution^11^, with notable examples including reduced genome size in birds with more aerial lifestyles^2^, accelerated evolution of regulatory mechanisms upon the loss of flight^12^, and major genomic changes associated with bone structure and song complexity^1,4,5^. However, it remains a challenge to disentangle the roles of life-history traits in driving genome evolution, and to identify any particular genes that have a strong association with the diversification of avian phenotypes^11,13^. The increased availability of whole genomes for large numbers of bird species^14^ offers an unprecedented opportunity to explore the association between life-history traits and genomic signals across evolutionary time^15^. In particular, dated phylogenetic trees provide a comparative framework that allows the detailed analysis of genomic evolutionary rates.

Studies of the rates of genomic change across taxa have helped to explain various macroevolutionary phenomena, and have the potential to improve our understanding of the factors that drive evolutionary success. For instance, the explosive diversification in birds has been linked to higher rates of molecular evolution^16^. Evolutionary rates are also likely influenced by life-history traits such as generation length^17^, metabolic rate^18^, and polygyny^19^, demonstrating a close relationship between ecology, genome evolution, and macroevolutionary success.

Genomic comparative research has often focused on variation in rates across taxa, while assuming that these patterns are consistent across all genomic regions^20^. Although among-lineage rate variation is an important form of heterogeneity in the molecular evolutionary process^21,22^, this focus overlooks any nuanced signals across the genome that might reveal the role of particular metabolic pathways or subsets of genes in driving evolutionary rates. In birds, for example, sex chromosomes and macrochromosomes appear to evolve more quickly than the rest of the genome^1,14^. This pattern possibly reflects the lower effective population size of sex chromosomes and the reduced gene density in large chromosomes. Similarly, accelerated rates might be apparent in the coding regions of songbird genomes, since complex vocalization is likely associated with dramatic genomic changes^5^. Whole-genome data from large numbers of avian taxa now offer a wealth of information about the molecular evolutionary process. However, these data pose a challenge for the methods that are commonly used to study evolutionary rate variation, which generally have limited scalability.

Here we identify the major axes of evolutionary rate variation across phylogenetic branches and genomic loci using principal components analysis of molecular rate estimates. This method is the first to maintain the rates data structure while modelling main axes of variation and allowing formal tests of influential taxa and loci at each axis. We refer to this approach as ‘evolutionary rate decomposition’ (Extended Data Fig. 1). We implement this method across the data set of 218 *de novo* genomes produced by the B10K consortium^14^ sampled for almost every extant avian family (87%), allowing us to identify axes of genes that have covarying rates within subsets of lineages^23^. Using a comprehensive data set of traits at the species level, we identify the major correlates of evolutionary rates across families. We also leverage estimates of rates of nonsynonymous substitutions (*d*_N_) and synonymous substitutions (*d*_S_) to gain insights into the contribution of mutation rate, selection, and population size to rate variation^24^. Decomposition of evolutionary rates into major axes of variation across lineages allows us to present a comprehensive description of the dominant drivers of evolutionary speed at the genomic scale.

### Clutch size predicts genome-wide mutation rate

To examine the relationship between biological traits and molecular evolution in birds, we analysed a comprehensive set of 29 life-history, morphological, ecological, geographical, and environmental traits (Supplementary File 1; Methods). We tested the association between these traits and average family-level rates across four measures of molecular evolution: *d*_N_, *d*_S_, *d*_N_/*d*_S_ (*ω*), and rates in intergenic (IG) regions. These measures have the potential to provide complementary insights into the evolutionary process: *d*_N_ is influenced by the mutation rate, population size, and selection; *d*_S_ is influenced by the mutation rate; and their ratio (*ω*) is influenced by selection and population size. Genome-wide variation in *ω* is expected to be driven by population size. Rates in intergenic regions of the genome are expected to reflect the mutation rate. We first used simple phylogenetic regression to test the hypothesis that each individual trait explains each molecular rate metric. Next, we used multiple regressions to test whether individually significant traits as partial terms explain each molecular rate metric.

Clutch size showed a significant positive association with mean *d*_N_, mean *d*_S_, and mean rates in intergenic regions in simple and multiple regressions (Fig. 1a, multiple regression *p_d_*_N_ = 0.048, *R*^2^*_d_*_N_ = 0.246, *p_d_*_S_ < 0.001, *R*^2^*_d_*_S_ = 0.362, *p*_IG_ = 0.004, *R*^2^_IG_ = 0.399). Generation length was the only other variable associated with molecular rates in multiple regression, showing a negative relationship with rates in intergenic regions (Fig. 1b, multiple regression *p*_IG_ = 0.029, *R*^2^_IG_ = 0.399). These results suggest a role for clutch size and generation length in driving mutation rates across deep time scales, complementing recent results that identified both variables as predictors of instantaneous mutation rates in parent-offspring trios^25^. Clutch size is a life-history trait that is likely to play an important role in adaptation, given its association with a wide range of other species characteristics, including growth rate, egg size, and longevity, and geographic factors such as insularity and latitude^26^. Meanwhile, animals with short generation times tend to copy their genomes more frequently per unit of time, whereas those with long generation times are expected to place greater investment in DNA repair^17,25^. Also, longer-lived animals tend to have smaller clutch sizes, such that the two factors are expected to contribute jointly to lower rates. Clutch size might offer greater explanatory power for molecular rate variation than other variables because of its impact on the number of viable genomic replications per generation. For instance, larger clutch sizes are associated with greater numbers of viable copies of the genome, increasing the opportunity for mutations to be transmitted to future generations^27^. Alternatively, the expected greater parental care in small clutches could lead to reduced mutation rates, driven by reduced exposure to mutagens in the germ line^28^. However, this latter hypothesis has not been described or tested in detail.

**Fig. 1.**
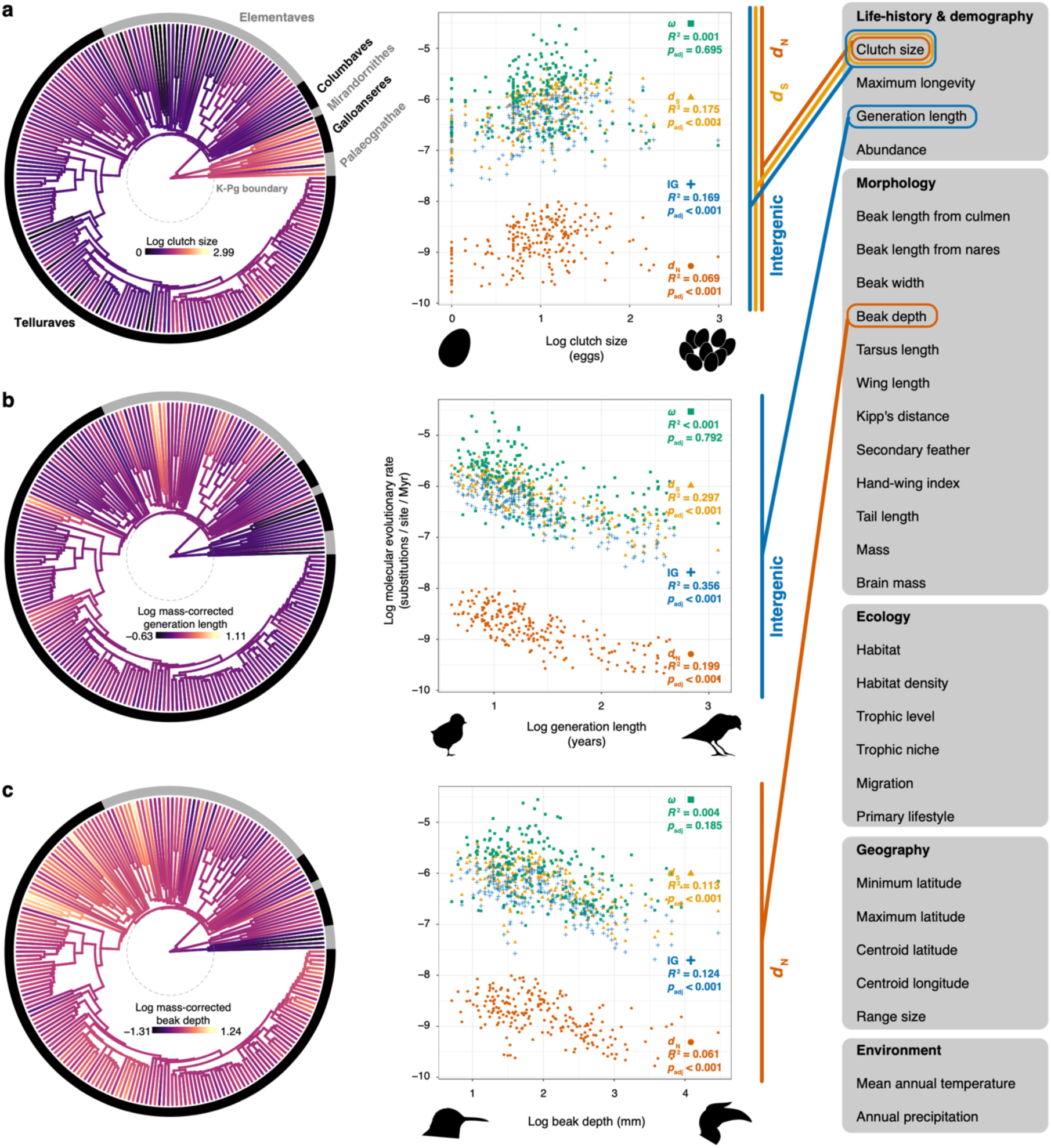
Association between genome-wide family-level molecular rates and avian traits. Regression *p*-values are adjusted for false-discovery rates, and together with *R*^2^ values correspond to results of simple regression models. The terms were qualitatively identical in multiple regression models, in which traits as partial terms explained *d*_N_, *d*_S_, and rates in intergenic regions. Branch colours show uncorrected log mean clutch size per family. **a** Clutch size significantly predicted the most metrics of genome-wide molecular rates, while **b** generation length and **c** beak depth were also important predictors of rates at intergenic regions and *d*_N_, respectively.

An additional finding is the negative association between beak depth and genome-wide *d*_N_ (Fig. 1c, multiple regression *p_d_*_N_ = 0.011, *R*^2^*_d_*_N_ = 0.246). Beak shape has previously been linked with adaptation in birds, not only as an intrinsic predictor of diet^29^, but also a predictor of adaptation to climate^30,31^, trophic interactions^32,33^, and interspecific competition^34,35^. Our results are consistent with beak shape having wide-ranging and thus genome-wide adaptive importance in birds. Among shallow-beaked taxa with higher *d*_N_ rates are diverse families of insectivores, nectarivores, and dietary specialists, such as warblers, swallows, swifts, tyrants, treecreepers, nightjars, and hummingbirds. At the opposite end, deep-beaked taxa include several families that are less diverse, often with relatively restricted geographic ranges and carnivorous or generalist habits, such as the shoebills, hornbills, secretary birds, ostriches, parrots, and vultures. The ecological differences among families with differing beak depth suggest that taxa with shallow beaks might have greater trophic and climatic flexibility, explaining their higher rates of genome-wide adaptive evolution.

When excluding flightless palaeognath taxa with extreme values of wing shape (Struthionidae and Rheidae), we additionally found tarsus length and brain mass to have a negative association with genome-wide *d*_N_ (multiple regression *p*_tarsus length_ = 0.038, *p*_brain mass_ = 0.008, *R*^2^_dN_ = 0.278, Extended Data Fig. 2). Tarsus length is also linked to intergenic region rates in these analyses (multiple regression *p*_tarsus length_ = 0.044, *R*^2^_IG_ = 0.417). Short tarsi are often associated with perching and prey clawing, while long tarsi often occur in ground-dwelling birds. Thus, this result might be consistent with genomically widespread adaptations to flight-intensive lifestyles associated with shorter tarsi. Another possible link between tarsus length and brain mass and genome-wide *d*_N_ is via population size. For example, taxa with large brains like corvids are generalists in terms of size, morphology, lifestyle, and foraging, which is likely to allow them to maintain large population sizes. Taxa with smaller brains and longer tarsi such as cariamids and flamingos are often ground-dwelling, and their reduced intensity of flight might lead to low connectivity among populations^36^. Nonetheless, traits associated with flight habit, like the hand-wing index, were not significant predictors of molecular rates in our data.

Body size is correlated with several life-history traits and is often found to be associated with molecular evolution in vertebrates, yet in birds we have found no evidence that body size predicts variation in molecular rates. Body size in birds apparently underwent substantial reductions immediately after extinction events including the Cretaceous–Palaeogene (K–Pg) transition, in what is known as the “Lilliput effect”^37,38^. Large body sizes are predicted to be associated with greater number of cells under pressure to reduce somatic mutation and damage, particularly when body size is associated with longevity^17^. Our data suggest that body size is not a dominant driver of genome-wide evolutionary rate, but give more prominent roles to clutch size and other factors.

The lack of support for any association of genome-wide *ω* with body size or any other trait points to a limited role of variations in population sizes on avian molecular evolution. Population sizes are believed to have increased rapidly immediately after the K–Pg transition, during a period of newly available ecological niches. If this were the case, then we might also expect lower *ω* in smaller-sized avian taxa, consistent with the Lilliput effect, due to the more efficient removal of slightly deleterious nonsynonymous mutations^24^. One explanation for the lack of this signal in our data is that the effects of population-size fluctuations can be seen primarily on shallower timescales rather than in deep lineages where the signal has eroded. Alternatively, our finding of variation in *ω* across axes of variation and across chromosomes suggests that the impact of population size might be uneven across the genome (Stiller et al. submitted). Lastly, incomplete lineage sorting and reticulate evolution can also degrade the signal of ancestral population size^39^. Future examinations of these questions at finer taxonomic levels will be able to show a more nuanced picture, for instance by extending estimates of population size dynamics^40^ into deeper timescales.

### Major changes in evolutionary rate across lineages

Rate decomposition is a powerful approach to analysis of molecular rates that allows a dissection of the subsets of lineages and of genes that have had an unusually large impact on evolutionary rate variation^23^ (Extended Data Fig. 1). By using principal components analysis, the method uses unsupervised learning for modelling the main axes of variation in rates, while permutation can be used to test axes and each taxon in each axis for their contribution to variation in rates. Out of 433 principal components in each avian molecular data set, permutation tests consistently showed that <5% of components explained significant amounts of variation in rates (*p* < 0.01, 8 principal components in *d*_N_, 10 in *d*_S_, 7 in *ω*, and 16 in intergenic regions).

The first principal component explained large portions of variation in *d*_N_ (35%), *ω* (38%), and rates in intergenic regions (24%), and the lineages with the greater loadings on this component were consistently terminal branches (Fig. 2, Extended Data Fig. 3). This indicates that large numbers of family lineages share the same pattern of rate variation across loci. Such a result likely reflects the dominating effect of gene-specific rates rather than the signals from subsets of phylogenetic lineages^21^. Since this result of terminal branches dominating rate heterogeneity is observed in *d*_N_ and *ω*, but not *d*_S_, it is likely to reflect widespread variation in genome-wide selection constraints or population sizes within families. Denser taxon sampling will enable further disentanglement of the drivers of rate variation, such as any concomitant climate-driven fluctuations in population size across families^41^. Alternatively, the strong influence of terminal branches might indicate the difficulty in recovering signals of selection or population size fluctuation at deeper timescales.

**Fig. 2.**
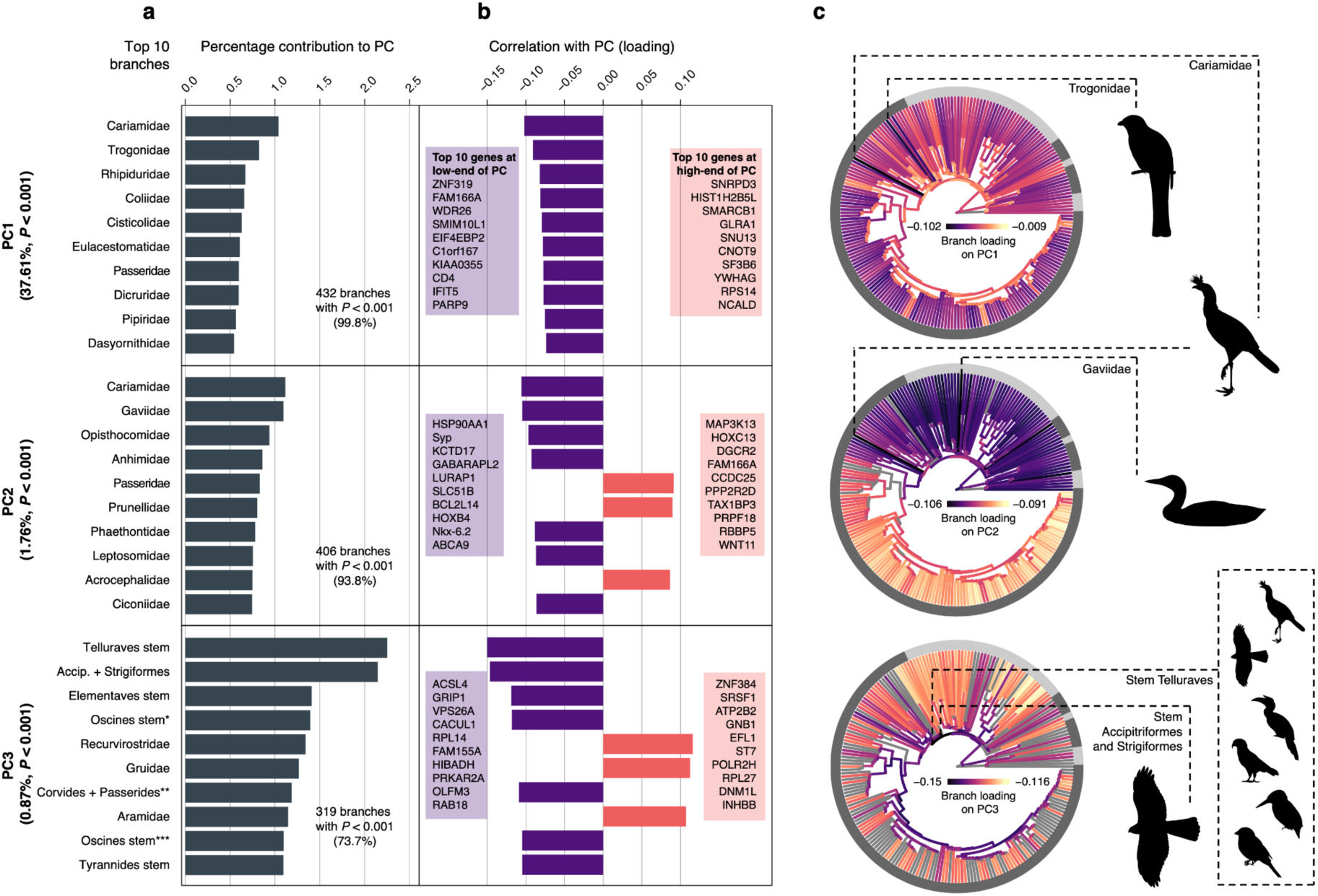
Contribution to variance and loading across taxa on principal components of *ω* across loci. The rates matrix that undergoes decomposition includes branches as columns and loci as rows, such that principal components describe the variation of locus rates across axes that can be explained by different sets of taxa (Extended Data Fig. 1). **a** Taxon contributions for the top ten taxa in the top three principal components. The principal components and taxa shown here explain significantly more variation than expected under permutation (*p* < 0.01). **b** Branch loadings indicate the correlation between taxa and locus rates in each principal component. **c** Branches are coloured by loadings, where grey branches are those that did not have significantly greater explanatory power than expected under permutation. The outer ring indicates the major taxonomic groups named in Fig. 1, while dashed lines indicate the two branches with the greatest loading in each principal component. According to the taxonomy of passerines^43^: * indicates the stem of the Oscines minus the grouping of Menuridae and Atrichornithidae; ** the stem of infraorders Corvides and Passerides, plus the grouping of families Orthonychidae and Pomatostomidae; *** the stem of the stem of the Oscines minus Menuridae, Atrichornithidae, Climacteridae, and Ptilonorhynchidae.

The other significant principal components of *d*_N_ and *ω* highlighted the stem branches leading to Telluraves and Elementaves, as well as several deep branches in the passerine clade, as being dominant drivers of variance in molecular rates (Fig. 2). The prominent role of these early neoavian branches in explaining variation in molecular rates points towards major ecological and adaptive changes across birds occurring immediately after the K–Pg transition. This is consistent with evidence of changes in a range of traits such as body size and brain mass at that time^38,42^. The major principal components of *d*_S_ variation also point to the same branches (stems of Telluraves and Elementaves) as having the greatest explanatory power, consistent with genome-wide increases in mutation rate soon after the K–Pg transition.

### Extensive variation in avian chromosomal rates

Evolutionary rates show considerable variation across chromosomes, with macrochromosomes frequently having significantly lower rates than expected under permutation of loci (Fig. 3a). Similarly, microchromosomes frequently have significantly higher rates than expected under permutation. These results are consistent with previous findings of greater rates of intrachromosomal rearrangement and recombination in micro-than in macrochromosomes^44^(Stiller et al. submitted). Microchromosomes have also been found to contain greater amounts of simple repeats and GC-rich hotspots of double-strand breakage. Therefore, the increased GC content that arises from recombination is a possible source of DNA damage, and a likely cause of elevated mutation rate in microchromosomes.

**Fig. 3.**
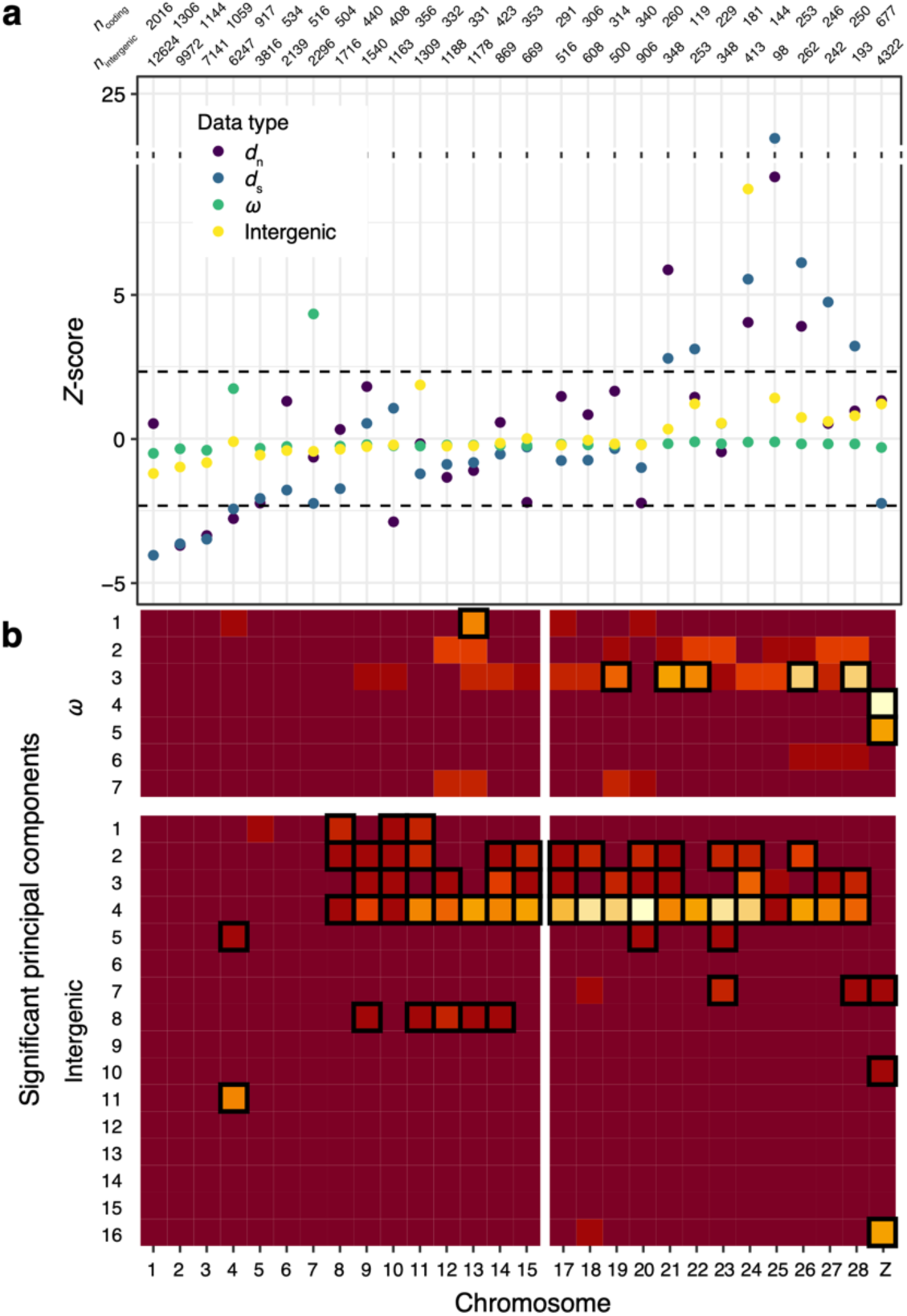
Avian mean chromosome rates and chromosomal representation across principal components of rate decomposition analyses. **a** Each chromosome was assessed for its mean rate being different from that expected under genome-wide permutation. **b** Loci at the extremes of principal components were tested for having an overrepresentation of each chromosome. Light colours indicate increasing overrepresentation of a chromosome (rows) at a particular principal component (columns). Boxed cells indicate chromosomes with a significantly greater portion of genes at the extremes of a principal component (*p* < 0.01). Chromosome 16 was excluded from all analyses due to a small sample size of loci.

Rate decomposition further revealed the subsets of lineages in which chromosomes showed extremes in rates. The extreme values of principal components of *ω* and intergenic regions included an overrepresentation of microchromosome loci (Fig. 3b). This finding on *ω* rates is consistent with widespread selective constraints on microchromosomes immediately after the K–Pg transition. This result is also consistent with the critical role that has been proposed for microchromosomes in harbouring housekeeping genes. Intergenic regions similarly show extreme rates occurring primarily in microchromosomes. However, the principal components of these extreme rates are explained by early passerine branches (Extended Data Fig. 3).

The overrepresentation of microchromosomes in the evolutionary rate shifts along early neoavian branches suggests that they have had a greater influence on avian evolution than macrochromosomes. This result is also consistent with evidence that microchromosomes are highly acetylated powerhouses of gene expression, and the building blocks of larger chromosomes^45^. Meanwhile, macrochromosomes are less resistant to major rearrangements and their large size is a consequence of the fusion of microchromosomes that are less transcriptionally active^44^. Evolutionary rates are likely to have varied the most in microchromosome regions undergoing intense selection, fundamental for development and adaptation. Since our results of molecular rates focus on evolution at the nucleotide level, they suggest that even under high mutational pressure synteny can be conserved across deep timescales in microchromosomes, reflecting a balance between high nucleotide mutation rates even under severe selective constraints.

### Early shifts in meiotic, cardiac, and gene regulation machineries

To investigate the functional groups of genes associated with the most prominent branches driving rate variation, we examined loci with extreme values at principal components. To understand the ecological implications of the genes at extremes of rate axes, we also examined whether traits of families were associated with the proportion of variance explained (i.e., loadings on principal components). Gene set enrichment analysis was performed for the 20% of loci at the extremes of principal components of *ω* (Extended Data Fig. 1), focusing on those components that explained significantly greater heterogeneity than expected under permutation (*p* < 0.01). Along the first principal component of *ω*, enrichment was found for the ribosome gene term at high values of the axis (Supplementary File 2). Since this axis is negatively linked with rates on terminal branches, the result indicates a marked slowdown in ribosomal evolutionary rates across many present-day avian families.

The loadings at the third principal component of *ω* showed a significant positive relationship with tarsus length (Fig. 4a, *p_adjusted_* < 0.001, *R*^2^ = 0.064), suggesting that genes with greater values on this axis had accelerated rates in taxa with long tarsi. This was the only significant association found in both simple and multiple regressions between family loadings and any trait variable across principal components (Supplementary File 3). This principal component (PC3 of *ω*) was also one of the few where large numbers of taxa still contribute to variation significantly more than expected under permutation. These terminal branches leading to birds with long tarsi represent a diverse and ancient set of avian families including including plovers (Charadriidae), cranes (Gruidae), several raptors (Accipitridae, Pandionidae, Sagittariidae), and some passerines (Hyliidae, Cardinalidae, Thraupidae).

**Fig. 4.**
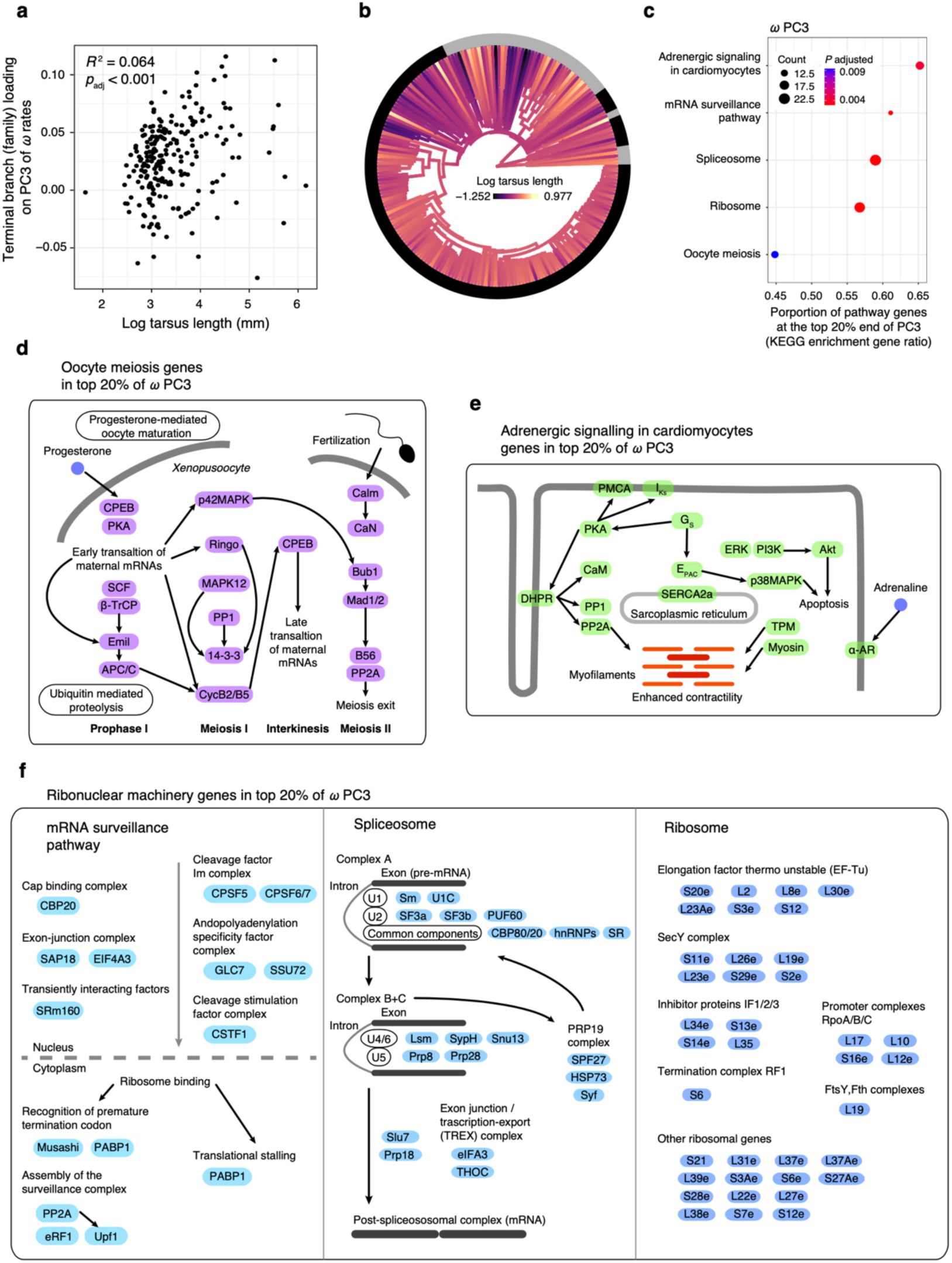
Trait regression and gene enrichment associated with PC3 of *ω* decomposition. **a** Regression *p*-value and *R*^2^ are shown for the simple regression among family-level (terminal branches) loadings on *ω* PC3 against log mean tarsus length. This variable was also uniquely significant in multiple regression involving other traits as partial terms. **b** The phylogenetic distribution of tarsus length shows the increase in diversity in this trait immediately after the K–Pg transition. Genes with extreme values of the third principal component of *ω* were enriched for **c** oocyte meiosis, putatively linked to DNA repair mechanisms, and **d** adrenergic signalling in cardiomyocytes, linked with heart function. **e** Additional enriched terms included the ribonuclear machineries for splicing, mRNA surveillance, and ribosomal activity. Only the genes inside the extreme values of PC3 of *ω* are shown, such that their metabolic links are approximate.

One distinctive feature of tarsus length is its very rapid increase in diversity immediately following the K–Pg transition (Fig. 4b). The broad range of adaptations and lifestyles associated with tarsus length include predatory habits such as those found in many waterbirds and raptors, and the ability to forage while perching as seen in many passerines. The rapid early evolution of this trait suggests strong positive selection that can be explained by the enriched gene terms at the ends of PC3 of *ω*. In addition to corresponding to accelerated molecular rates along terminal branches of birds with long tarsi, these terms indicate accelerated rates in some of the branches with the greatest loadings on this axis, including stem Telluraves, stem Elementaves, and early passerine branches (Fig. 2).

In addition to accelerated rates in taxa with long tarsi and decelerated rates in deep post-K–Pg branches, we found the top end of PC3 of *ω* to be enriched for five gene functional terms (Fig. 4c). One of these terms was oocyte meiosis (Fig. 4d), which is the primary pathway involved in the replication machinery of the germline and can be expected to have widespread effects across the genome through its impact in mutation rate, and hence estimates of *d*_S_. Interestingly, the principal components of *d*_S_ did not show enrichment for any gene term and instead consistently supported stem branches of Telluraves and of Elementaves as dominant drivers of evolutionary rate changes (Extended Data Fig. 3). One explanation is that in addition to a broad range of adaptive changes, avian genomes were also subject to changes in meiotic DNA repair mechanisms that led to increased mutation rates along those early branches. Support for this hypothesis includes the presence of genes that are directly involved in DNA repair among those in the enriched oocyte meiosis term, such as MAPK^46,47^. A further finding was the enrichment of genes coding for adrenergic signalling in cardiomyocytes (Fig. 4e). This pathway is associated with cardiac performance^48,49^ and suggests fast evolution in heart function in old families with long tarsi, and was presumably involved in allowing early avian lineages to sustain intense flight habits.

Additional enriched pathways included the combination of the spliceosome, mRNA surveillance, and the ribosome (Fig. 4f). In combination, these terms form the backbone of the ribonuclear cellular machinery that regulates the quality control and translation of gene products. The evolution of ribonuclear machineries has been linked to protection against deleterious insertions and mutation^50–52^. The change in *ω* in ribonuclear machineries in a broad range of taxa with long tarsi suggests that genomic changes contributed to increased efficiency or novelty in gene expression and regulation. Ribonuclear machinery evolution in some of the oldest families with long tarsi may have played a role in the colonization of a vast diversity of habitats, contributing to the rapid radiations of these groups after the K–Pg transition. The marked deceleration in *ω* in stem Telluraves and stem Elementaves might reflect large population sizes that were probably seen after the K–Pg transition. It is likely that there was widespread competition among early avian lineages at this time, and changes in fundamental ribonuclear machineries in early Neoaves could have conferred a competitive advantage, providing some key steps towards the explosive diversifications that followed. Taken together, these results indicate that it was during the short period after the K–Pg transition that a dramatic increase in novel adaptations took place in stem neoavians, and that these changes were likely facilitated by evolution in ribonuclear machineries.

### Conclusions

Modern molecular clock theory can be used in combination with large numbers of whole genomes to disentangle processes of mutation and selection over deep time scales. We have demonstrated that identifying the dominant axes of rate variation provides useful insights into the relationships between biological diversity and genomic evolution. To achieve this, we performed a series of genomic comparative analyses that revolved around the decomposition of evolutionary rates across regions of the genome and across phylogenetic branches. Our genome-scale molecular rate analyses show that clutch size, generation length, and beak depth – traits long considered important in avian life history and ecology – are key drivers of genome evolution in birds. Our decomposition of evolutionary rates also revealed the importance of meiotic, cardiac, and ribonuclear machinery evolution along some of the deep branches in the avian tree. These rate shifts took place primarily in microchromosomes and in taxa with longer tarsi – a trait linked with terrestrial locomotion. Collectively, our analyses indicate that meiotic and ribonuclear machineries and microchromosomes, with crucial roles in DNA replication, gene usage, and housekeeping, underwent rapid evolution that likely contributed to the explosion of avian diversity filling vacant niches soon after the K–Pg transition. We expect that increasing numbers of genomes gathered from large numbers of taxa will continue to offer valuable insights into the primary evolutionary processes shaping phenotypic variation across the avian tree of life.

## Data availability

Genomic assemblies and annotations form avian families orginally from Feng *et al.*^14^ are available in the NCBI SRA and GenBank under accession PRJNA545868. Gene trees and species tree data are available via https://sid.erda.dk/cgi-sid/ls.py?share_id=hVS3naBtJ6 while trait data and analysis scripts are available at https://github.com/duchene/avian_family_rates.git and will receive a permanent DOI upon acceptance of the article. This latter repository also contains copies of supplementary files and instructions on extended visualisation of results.

## Supplementary information

**Supplementary File 1**: Table describing avian trait variables and their sources.

**Supplementary File 2**: Enriched pathways and genes involved.

**Supplementary File 3**: Tables of phylogenetic regression data and results.

## Acknowledgements

This work was supported by grants from the Carlsbergfondet of Denmark (grant CF18-0223) and a European Research Executive Agency Marie Sklodowska-Curie Action (H2020-MSCA-IF-2019-883832) to D.A.D., and the Danish National Research Fund ‘Center for Evolutionary Hologenomics’ award to M.T.P.G. (DNRF143). S.F was supported by a National Natural Science Foundation of China grant (number 32170626). The international collaboration was supported by a Partnership Collaboration Award from the University of Sydney and University of Copenhagen to D.A.D., M.T.P.G., G.Z., and S.Y.W.H. Images of metabolic pathways are interpretations from the R package pathview. Silhouettes of birds and bird parts are originals or modifications from phylopic.org dedicated to the public domain.

## Author contributions

D.A.D., S.Y.W.H., A-A.C., J.A.T., M.T.P.G., G.Z., and S.F. conceived and designed the study. D.A.D., A-A.C., J.S., J.A.T., and J.Y. contributed to data collection and preparation. D.A.D. and A-A.C. performed genomic analyses and regression analyses. D.A.D., S.Y.W.H., A-A.C., M.I-C., J.A.T., G.Z., and S.F. contributed to the data interpretation. D.A.D., S.Y.W.H. and M.I-C. wrote the manuscript with input from all co-authors.

## Methods

Family-level phylogenomic data for avian families as published by Feng *et al.*^14^ were collected from the database of the B10K consortium. These included 63,430 evenly spaced intergenic loci each with 1 kbp in length, as well as the coding regions of 15,093 orthologous genes. For each data type, we selected the representative of each avian family for which the greatest nucleotide completeness was available, providing a sample of 218 sampled tips. To make reliable estimates of branch lengths in expected synonymous (*d*_S_) and non-synonymous (*d*_N_) substitutions, coding regions were filtered to exclude codons where any of the three positions was missing for >50% of the taxa, and where the most common amino acid occurred in <50% of taxa. In each region, we also excluded taxa for which <30% of the nucleotides were available. Regions with fewer than three taxa remaining after this step were also excluded from further analyses, producing a final data set with 63,174 intergenic and 14,849 coding regions.

To obtain molecular evolutionary rate estimates for each genomic region, we used phylogenetic inferences of tree and branch lengths after forcing the tree topology. We forced the trees for each region to conform to the B10K family-level relationships, using the species tree inferred from intergenic regions under the multi-species coalescent (Stiller et al. submitted). Inferences of phylogenetic branch lengths from intergenic regions were performed using the best-fitting model from the GTR+F+R^53^ family in IQ-TREE v2.1.2^54^. Coding regions were used for inference of *d*_N_ and *d*_S_ branch lengths. We first estimated distance matrices for *d*_N_ and *d*_S_ using maximum likelihood in PAML using the models HKY+Γ and F3×4 for nucleotide and codon substitutions, respectively^55^. For each gene, maximum-likelihood *d*_N_ and *d*_S_ distance matrices were then transformed to the lengths of branches in the species tree using ERaBLE^56^. We then used ClockstaRX^23^ to extract branch lengths for each locus at each species tree branch, which we then converted into rates using the family-level time-tree estimate. To avoid adding error due to lack of signal in the molecular data, we excluded branches with length estimates <5×10^-6^ substitutions/site. Analyses where each locus has a free topology are also possible in ClockstaRX, but led to a vast loss of data, including the majority of genes at deep branches, as expected in ancient systems with large amounts of phylogenetic error and incomplete lineage sorting. These analyses were not considered further.

To explore whether specific species traits explained molecular rates, we collected trait data from a series of databases and primary literature (see Supplementary File 2 for a description of each variable). We collected a total of 29 traits with complete or near-complete sampling across bird species, including 4 life-history and demography traits (clutch size, longevity, generation length, and abundance), 12 morphological traits (beak length from culment, beak length from nares, beak width, beak depth, tarsus length, wing length, Kipp’s distance, secondary feather length, hand-wing index, tail length, body mass, and brain mass), 6 ecological traits (habitat, habitat density, trophic level, trophic niche, migration, and primary lifestyle), 5 geographic traits (minimum, maximum and centroid latitude, centroid longitude, and range size), and 2 environmental traits (mean annual temperature, annual precipitation). The family mean and mode were used for continuous and discrete traits, respectively. We first tested whether each individual trait was associated with the estimates of mean molecular rates in each family (in all *d*_N_, *d*_S_, *ω*, and intergenic regions), using phylogenetic regression as implemented in the package phylolm^57^. We then ran multiple phylogenetic regression, where all the significant explanatory traits from the individual regressions were added as explanatory variables as partial terms, testing whether significant variables explained molecular rates above the effect of other traits. All *p*-values were corrected for multiple comparisons using false discovery rates. Trait data were visualised across branches using fast maximum-likelihood ancestral state reconstruction as implemented in phytools^58^. Phylogenetic data were processed with the support of phangorn^59^.

We tested whether the mean molecular rate of each chromosome was different from the mean expectation from loci across the genome. To do this, we performed permutations per chromosome, involving a random draw of 1000 sets of loci of identical size to the chromosome. We performed this permutation for each chromosome and calculated the *Z*-score of the mean empirical chromosome rate with respect to the permutation distribution.

We performed rate decomposition analyses to identify the lineages and genes that dominate molecular rate variation. This was done in ClockstaRX^23^ for each set of loci independently (*d*_N_, *d*_S_, *ω*, and intergenic regions). To allow computational efficiency when examining intergenic regions, we analysed three random samples of 10,000 loci, and confirmed that results were consistent across each sample. The basic method of decomposition collects rate estimates in a matrix containing each locus across rows and lineages in the species-tree estimate across columns. This data structure is then decomposed for identifying the main axes of variation using principal components analysis. The software performs a permutation of the data matrix to test each principal component and lineage loading for contributing more variation than expected from a random sample across the data^60^ (α = 0.01).

We explored whether lineage contribution to variance at rate axes was associated with any traits in our comprehensive set, such that a principal component might be driven by the ecology or life history of families. To do this, we first extracted the loadings of families (terminal branches) on each of the principal components that significantly explained variation in rates. We then proceeded to perform simple and multiple phylogenetic regressions where traits were tested to explain principal component loadings, as performed for mean genome-wide rate metrics.

We then explored whether principal components are associated with any particular metabolic pathway or gene function. We extracted the 20% of loci with the maximum and minimum values at the principal components that significantly explained variation in rates. We used gene set enrichment analysis to evaluate any overrepresentation of metabolic functions of gene products independently in loci with extreme high and low rates at each significant rate axis. Gene identities were inferred using the best blastn match^61^, and used as input for testing enrichment of KEGG terms using clusterProfiler^62^. Genes in pathways were visualised and interpreted using pathview^63^. Bonferroni correction was used for *p*-values of significant terms.

Overrepresentation of chromosomes in the 20% extremes of significant rate axes (principal components) was evaluated using binomial tests. The null hypothesis was therefore a Bernoulli experiment^64^ where the proportion of loci in a given chromosome is expected to be equal in the 20% maximum and minimum loci of each axis. This test was performed for each chromosome and each significant principal component, with *p*-values corrected using false-discovery rates within principal components.

## Extended Data Figures and Tables

**Extended Data Fig. 1.**
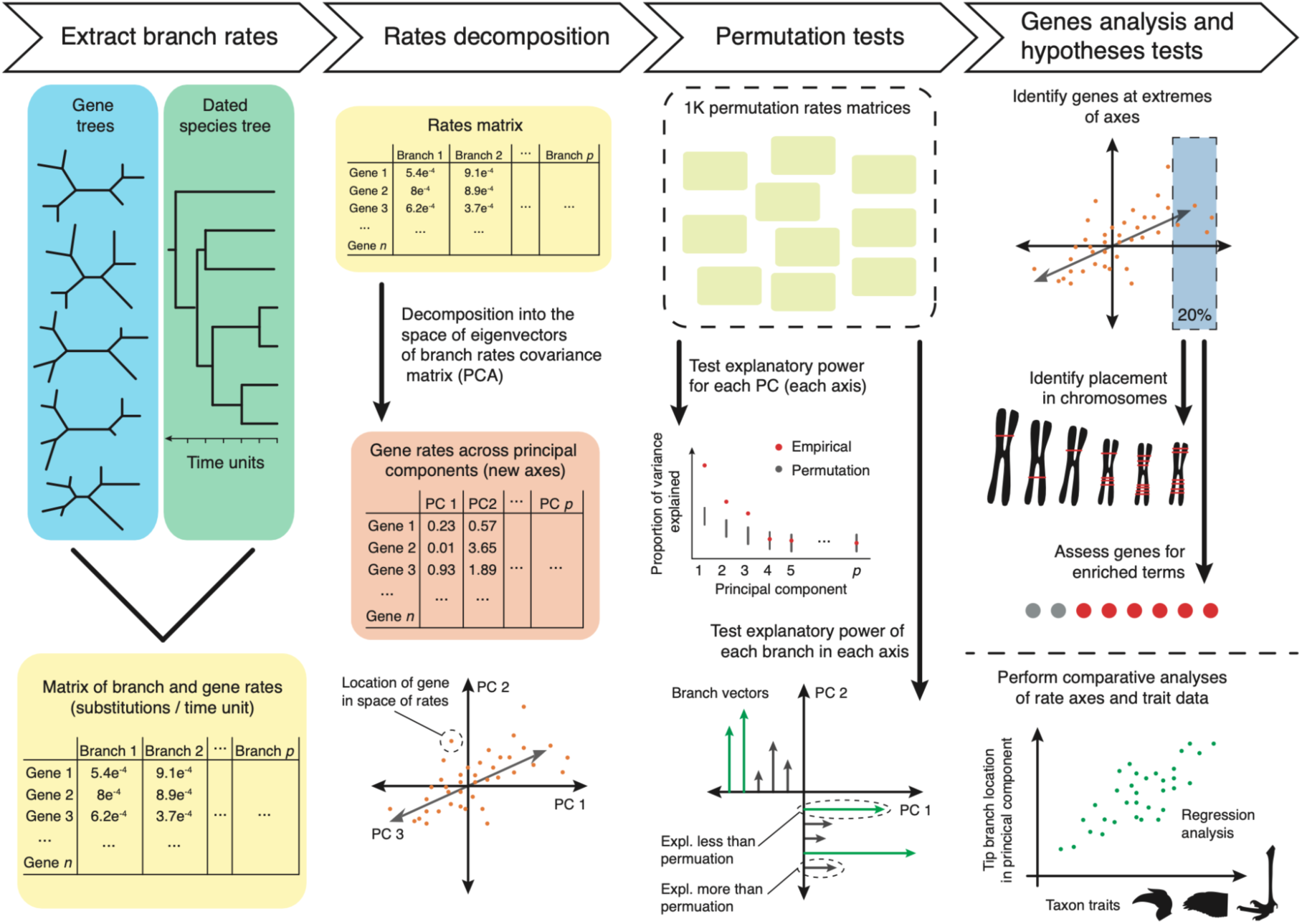
Overview of rate decomposition analyses. Data from gene trees and the species tree are the starting point, and are used as input in ClockstaRX to produce a rates matrix and perform rates decomposition and permutation analyses. The output can be used in a range of other downstream analyses, such as of gene enrichment and phylogenetic regression.

**Extended Data Fig. 2.**
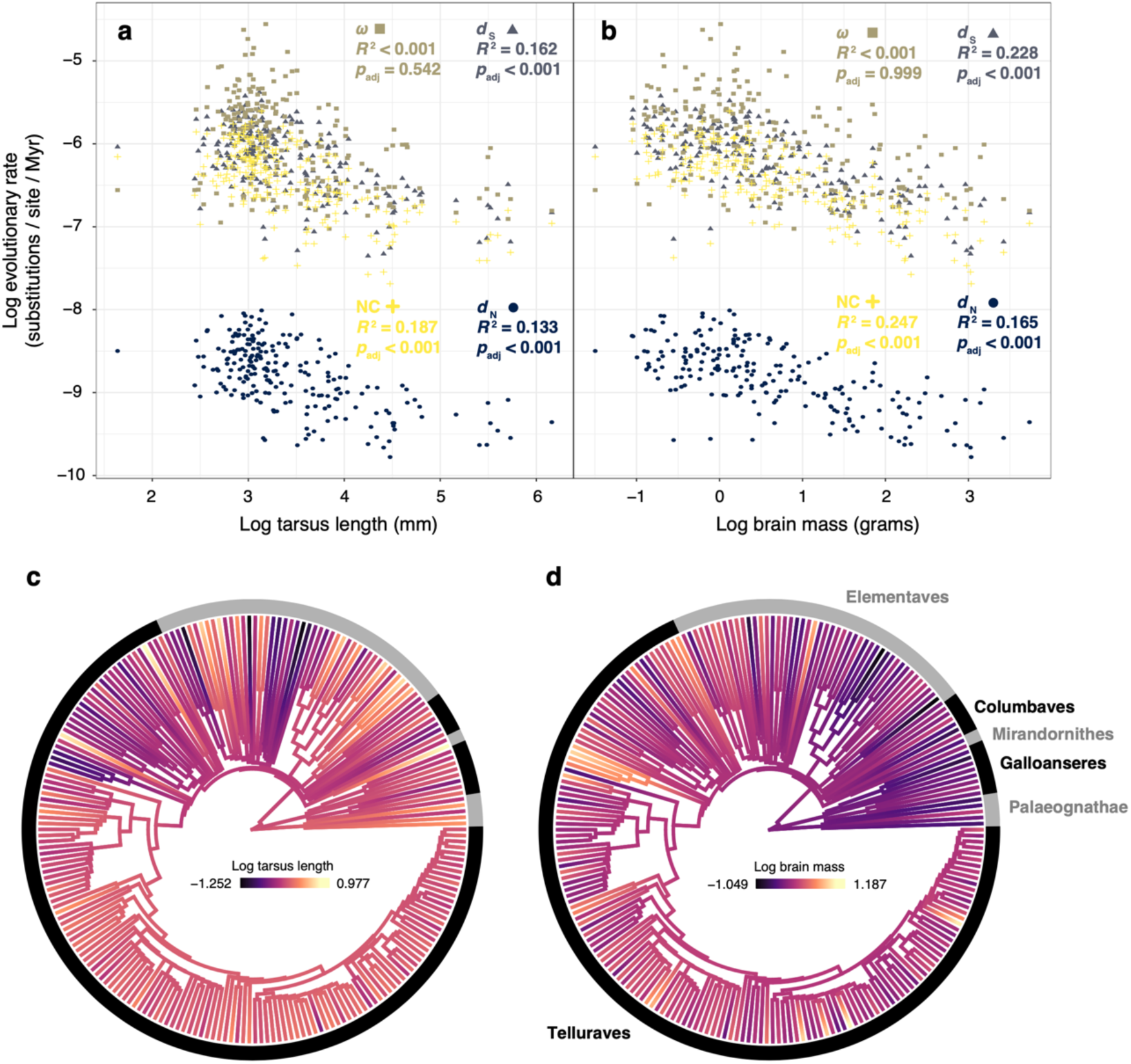
Significant phylogenetic regression results. After removal of extreme values of wing shape in flightless families (Struthionidae and Rheidae), tarsus length and brain mass emerged as significant in both simple and multiple phylogenetic regressions explaining *d*_N_ rates. The regression *p* and *R*^2^ values shown correspond to simple regression results adjusted for multiple comparisons using false discovery rates.

**Extended Data Fig. 3.**
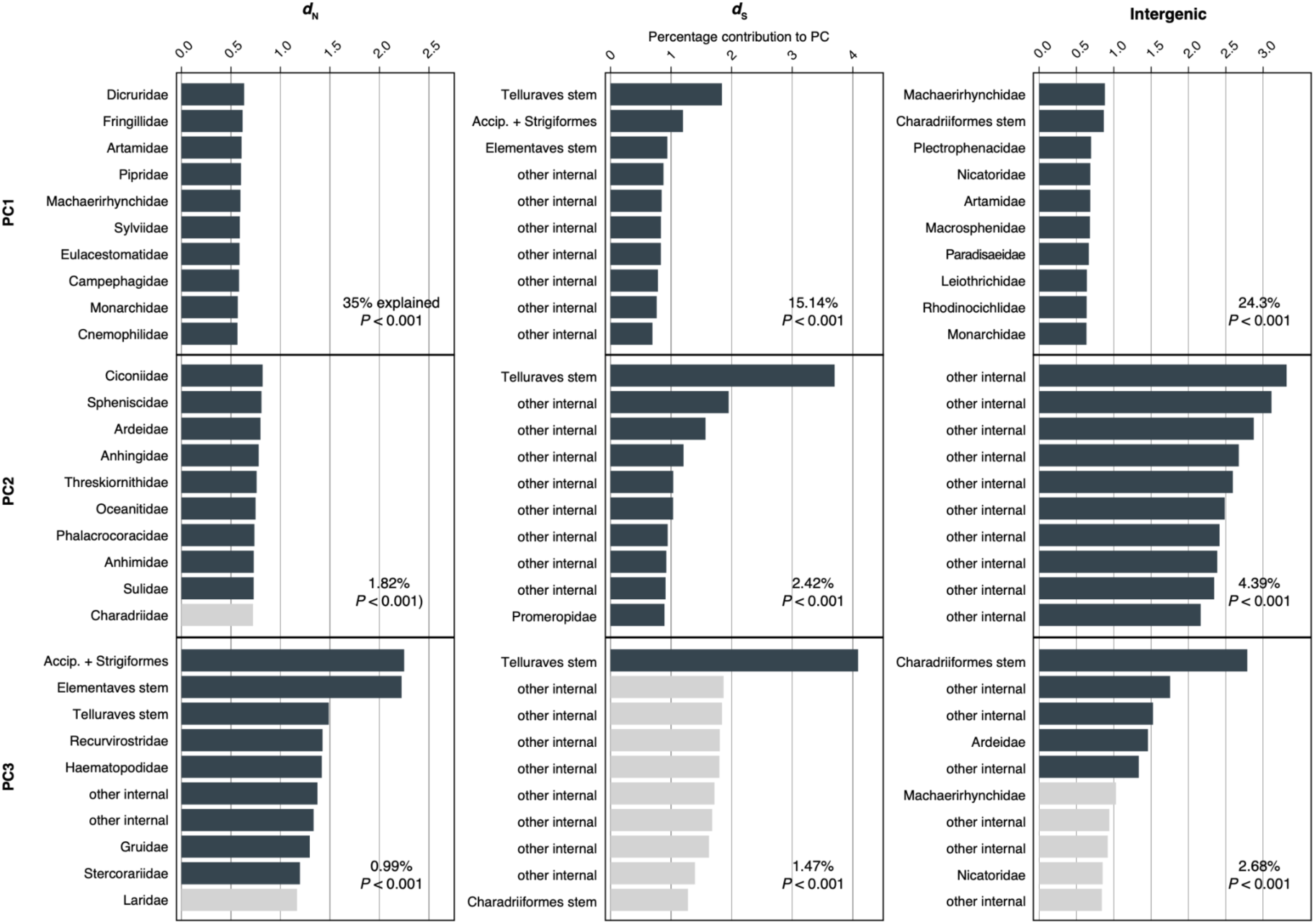
Taxon contribution to principal components. Rate decomposition for each molecular data set produces distinct results, yet consistently show a strong contribution of early neoavian branches (e.g., stem Telluraves). Results across all principal components can be found in the data repository listed in Data Availability.

**Extended Data Fig. 4.**
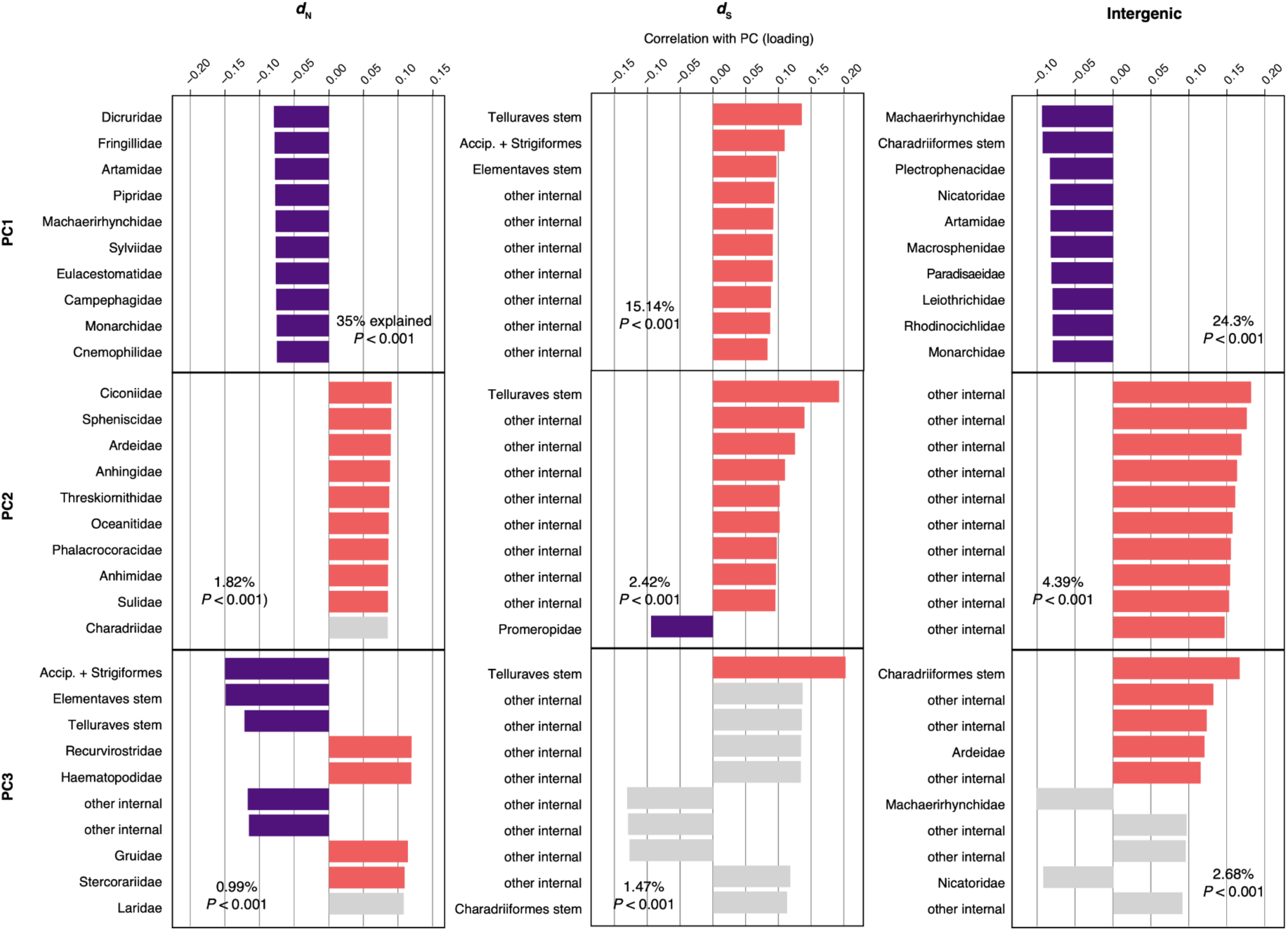
Correlation of taxa with each principal component. All branches had the same correlation sign with the first principal component of each data set, indicating that the principal component describes overall gene rate (rather than lineage-specific rates). Subsequent principal components contain more nuanced results, but frequently have large numbers of branches that do not explain variance significantly more than under permutation of the data.

**Extended Data Fig. 5.**
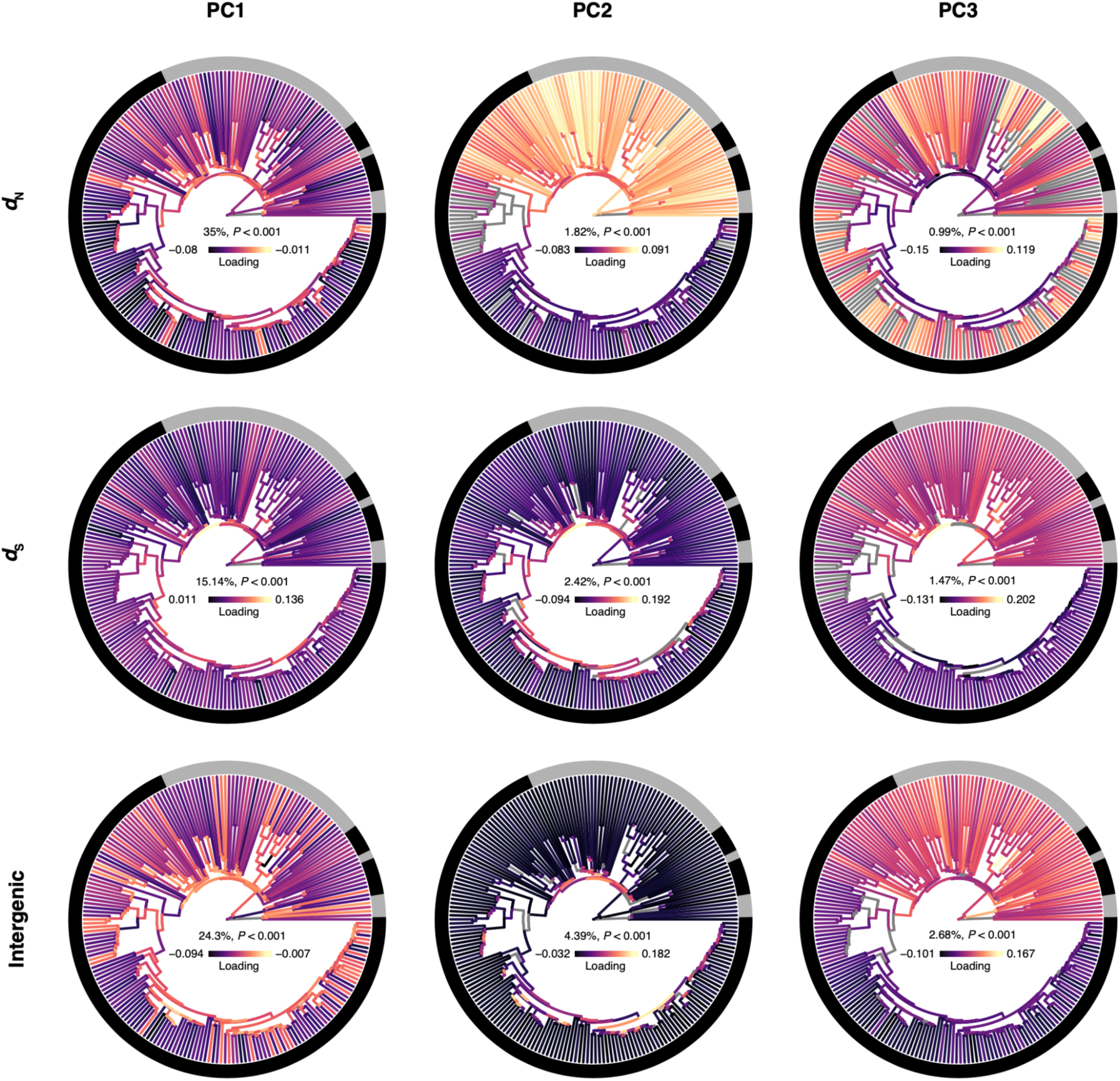
Branch loadings across principal components. Colours indicate the full range of loadings, yet first principal components consistently had the same loading sign across all branches. Branches shown in grey did not explain greater variance in the principal component than under permutation of the data.

## References

1. Zhang, G. et al. Comparative genomics reveals insights into avian genome evolution and adaptation. Science 346, 1311–1320 (2014).

2. Wright, N. A., Gregory, T. R. & Witt, C. C. Metabolic ‘engines’ of flight drive genome size reduction in birds. Proc. Biol. Sci. 281, 20132780 (2014).

3. Organ, C. L., Shedlock, A. M., Meade, A., Pagel, M. & Edwards, S. V. Origin of avian genome size and structure in non-avian dinosaurs. Nature 446, 180–184 (2007).

4. Whitney, O. et al. Core and region-enriched networks of behaviorally regulated genes and the singing genome. Science 346, 1256780 (2014).

5. Pfenning, A. R. et al. Convergent transcriptional specializations in the brains of humans and song-learning birds. Science 346, 1256846 (2014).

6. Mason, N. A. et al. Song evolution, speciation, and vocal learning in passerine birds. Evolution 71, 786–796 (2017).

7. Toomey, M. B. et al. High-density lipoprotein receptor SCARB1 is required for carotenoid coloration in birds. Proc. Natl. Acad. Sci. U. S. A. 114, 5219–5224 (2017).

8. Aguillon, S. M., Walsh, J. & Lovette, I. J. Extensive hybridization reveals multiple coloration genes underlying a complex plumage phenotype. Proc. Biol. Sci. 288, 20201805 (2021).

9. Abzhanov, A. et al. The calmodulin pathway and evolution of elongated beak morphology in Darwin’s finches. Nature 442, 563–567 (2006).

10. Yusuf, L. et al. Noncoding regions underpin avian bill shape diversification at macroevolutionary scales. Genome Res. 30, 553–565 (2020).

11. Bravo, G. A., Schmitt, C. J. & Edwards, S. V. What have we learned from the first 500 avian genomes? Annu. Rev. Ecol. Evol. Syst. 52, 611–639 (2021).

12. Sackton, T. B. et al. Convergent regulatory evolution and loss of flight in paleognathous birds. Science 364, 74–78 (2019).

13. Price-Waldman, R. & Stoddard, M. C. Avian coloration genetics: recent advances and emerging questions. J. Hered. 112, 395–416 (2021).

14. Feng, S. et al. Dense sampling of bird diversity increases power of comparative genomics. Nature 587, 252–257 (2020).

15. Smith, S. D., Pennell, M. W., Dunn, C. W. & Edwards, S. V. Phylogenetics is the new genetics (for most of biodiversity). Trends Ecol. Evol. 35, 415–425 (2020).

16. Lanfear, R., Ho, S. Y. W., Love, D. & Bromham, L. Mutation rate is linked to diversification in birds. Proc. Natl. Acad. Sci. U. S. A. 107, 20423–20428 (2010).

17. Bromham, L. Why do species vary in their rate of molecular evolution? Biol. Lett. 5, 401–404 (2009).

18. Montoya, P., Cadena, C. D., Claramunt, S. & Duchêne, D. A. Environmental niche and flight intensity are associated with molecular evolutionary rates in a large avian radiation. BMC Ecol. Evol. 22, 95 (2022).

19. Iglesias-Carrasco, M., Jennions, M. D., Ho, S. Y. W. & Duchêne, D. A. Sexual selection, body mass and molecular evolution interact to predict diversification in birds. Proc. Biol. Sci. 286, 20190172 (2019).

20. Lanfear, R., Welch, J. J. & Bromham, L. Watching the clock: studying variation in rates of molecular evolution between species. Trends Ecol. Evol. 25, 495–503 (2010).

21. Duchêne, D. A. et al. Linking branch lengths across sets of loci provides the highest statistical support for phylogenetic inference. Mol. Biol. Evol. 37, 1202–1210 (2020).

22. Snir, S., Wolf, Y. I. & Koonin, E. V. Universal pacemaker of genome evolution. PLoS Comput. Biol. 8, e1002785 (2012).

23. Duchêne, D. A., Duchêne, S., Stiller, J., Heller, R. & Ho, S. Y. W. ClockstaRX: testing molecular clock hypotheses with genomic data. bioRxiv 2023.02.02.526226 (2023) doi:10.1101/2023.02.02.526226.

24. Ohta, T. The nearly neutral theory of molecular evolution. Annu. Rev. Ecol. Syst. 23, 263–286 (1992).

25. Bergeron, L. A. et al. Evolution of the germline mutation rate across vertebrates. Nature 615, 285–291 (2023).

26. Jetz, W., Sekercioglu, C. H. & Böhning-Gaese, K. The worldwide variation in avian clutch size across species and space. PLOS Biol. 6, 2650–2657 (2008).

27. Brown, W. M., Prager, E. M., Wang, A. & Wilson, A. C. Mitochondrial DNA sequences of primates: tempo and mode of evolution. J. Mol. Evol. 18, 225–239 (1982).

28. Welch, J. J., Bininda-Emonds, O. R. P. & Bromham, L. Correlates of substitution rate variation in mammalian protein-coding sequences. BMC Evol. Biol. 8, 53 (2008).

29. Pigot, A. L. et al. Macroevolutionary convergence connects morphological form to ecological function in birds. Nat. Ecol. Evol. 4, 230–239 (2020).

30. Friedman, N. R., Harmáčková, L., Economo, E. P. & Remeš, V. Smaller beaks for colder winters: thermoregulation drives beak size evolution in Australasian songbirds. Evolution 71, 2120–2129 (2017).

31. Danner, R. M. & Greenberg, R. A critical season approach to Allen’s rule: bill size declines with winter temperature in a cold temperate environment. J. Biogeogr. 42, 114– 120 (2015).

32. Grant, P. R. & Rosemary Grant, B. 40 Years of Evolution: Darwin’s Finches on Daphne Major Island. (Princeton University Press, 2014).

33. Cattau, C. E., Fletcher, R. J., Jr, Kimball, R. T., Miller, C. W. & Kitchens, W. M. Rapid morphological change of a top predator with the invasion of a novel prey. Nat. Ecol. Evol. 2, 108–115 (2018).

34. Grant, P. R. & Grant, B. R. Evolution of character displacement in Darwin’s finches. Science 313, 224–226 (2006).

35. Freeman, B. G., Weeks, T., Schluter, D. & Tobias, J. A. The latitudinal gradient in rates of evolution for bird beaks, a species interaction trait. Ecol. Lett. 25, 635–646 (2022).

36. Claramunt, S., Derryberry, E. P., Remsen, J. V., Jr & Brumfield, R. T. High dispersal ability inhibits speciation in a continental radiation of passerine birds. Proc. Biol. Sci. 279, 1567–1574 (2012).

37. Harries, P. J. & Knorr, P. O. What does the ‘Lilliput effect’ mean? Palaeogeogr. Palaeoclimatol. Palaeoecol. 284, 4–10 (2009).

38. Berv, J. S. & Field, D. J. Genomic signature of an avian Lilliput effect across the K-Pg extinction. Syst. Biol. 67, 1–13 (2018).

39. Mendes, F. K. & Hahn, M. W. Gene tree discordance causes apparent substitution rate variation. Syst. Biol. 65, 711–721 (2016).

40. Germain, R. R. et al. Changes in the functional diversity of modern bird species over the last million years. Proc. Natl. Acad. Sci. U. S. A. 120, e2201945119 (2023).

41. Germain, R. R. et al. Species-specific traits mediate avian demographic responses under past climate change. Nat. Ecol. Evol. (2023) doi:10.1038/s41559-023-02055-3.

42. Ksepka, D. T. et al. Tempo and pattern of avian brain size evolution. Curr. Biol. 30, 2026–2036.e3 (2020).

43. Oliveros, C. H. et al. Earth history and the passerine superradiation. Proc. Natl. Acad. Sci. U. S. A. 116, 7916–7925 (2019).

44. Liu, J. et al. A new emu genome illuminates the evolution of genome configuration and nuclear architecture of avian chromosomes. Genome Res. 31, 497–511 (2021).

45. Waters, P. D. et al. Microchromosomes are building blocks of bird, reptile, and mammal chromosomes. Proc. Natl. Acad. Sci. U. S. A. 118, (2021).

46. Rezatabar, S. et al. RAS/MAPK signaling functions in oxidative stress, DNA damage response and cancer progression. J. Cell. Physiol. 234, 14951–14965 (2019).

47. Cannell, I. G. et al. p38 MAPK/MK2-mediated induction of miR-34c following DNA damage prevents Myc-dependent DNA replication. Proc. Natl. Acad. Sci. U. S. A. 107, 5375–5380 (2010).

48. Kim, J. et al. β-arrestin 1 regulates β2-adrenergic receptor-mediated skeletal muscle hypertrophy and contractility. Skelet. Muscle 8, 39 (2018).

49. Najafi, A., Sequeira, V., Kuster, D. W. D. & van der Velden, J. β-adrenergic receptor signalling and its functional consequences in the diseased heart. Eur. J. Clin. Invest. 46, 362–374 (2016).

50. Lynch, M. & Kewalramani, A. Messenger RNA surveillance and the evolutionary proliferation of introns. Mol. Biol. Evol. 20, 563–571 (2003).

51. Wolin, S. L. & Maquat, L. E. Cellular RNA surveillance in health and disease. Science 366, 822–827 (2019).

52. Singh, P., Saha, U., Paira, S. & Das, B. Nuclear mRNA surveillance mechanisms: function and links to human disease. J. Mol. Biol. 430, 1993–2013 (2018).

53. Kalyaanamoorthy, S., Minh, B. Q., Wong, T. K. F., von Haeseler, A. & Jermiin, L. S. ModelFinder: fast model selection for accurate phylogenetic estimates. Nat. Methods 14, 587–589 (2017).

54. Minh, B. Q. et al. IQ-TREE 2: new models and efficient methods for phylogenetic inference in the genomic era. Mol. Biol. Evol. 37, 1530–1534 (2020).

55. Yang, Z. PAML 4: phylogenetic analysis by maximum likelihood. Mol. Biol. Evol. 24, 1586–1591 (2007).

56. Binet, M., Gascuel, O., Scornavacca, C., Douzery, E. J. P. & Pardi, F. Fast and accurate branch lengths estimation for phylogenomic trees. BMC Bioinformatics 17, 23 (2016).

57. Ho, L. si T. & Ané, C. A linear-time algorithm for Gaussian and non-Gaussian trait evolution models. Syst. Biol. 63, 397–408 (2014).

58. Revell, L. J. phytools: an R package for phylogenetic comparative biology (and other things). Methods Ecol. Evol. 3, 217–223 (2012).

59. Schliep, K. P. phangorn: phylogenetic analysis in R. Bioinformatics 27, 592–593 (2010).

60. Björklund, M. Be careful with your principal components. Evolution 73, 2151–2158 (2019).

61. Camacho, C. et al. BLAST+: architecture and applications. BMC Bioinformatics 10, 421 (2009).

62. Wu, T. et al. clusterProfiler 4.0: A universal enrichment tool for interpreting omics data. Innovation (Camb) 2, 100141 (2021).

63. Luo, W. & Brouwer, C. Pathview: an R/Bioconductor package for pathway-based data integration and visualization. Bioinformatics 29, 1830–1831 (2013).

64. Clopper, C. J. & Pearson, E. S. The use of confidence or fiducial limits illustrated in the case of the binomial. Biometrika 26, 404 (1934).

